# IGF-1 from bone marrow Adipoq-lineage cells stimulates endocortical bone formation in mature female mice

**DOI:** 10.1101/2025.11.05.686616

**Authors:** Joshua C Bertels, Jasmin Koehnken Sawall, Brian Dulmovits, Xiaobin Liu, Ashley Phan, Xing Ji, Fangfang Song, Christopher Thom, Fanxin Long

## Abstract

Insulin-like growth factor 1 (IGF-1) is an anabolic signal promoting growth, differentiation and function of both embryonic and postnatal tissues. Both endocrine and paracrine functions of IGF-1 have been documented to regulate bone growth and bone marrow hematopoiesis. Local production of IGF-1 from various cell types may contribute differently to the overall bioactivity of IGF-1 in bone, but relevant sources and mechanisms are yet to be fully elucidated. Here we report that the Adipoq^+^ stromal cells are a notable source of IGF-1 in the bone marrow of postnatal mice. Deletion of IGF-1 with Adipoq-Cre diminished endocortical bone formation and cortical bone mass in mature female mice. On the other hand, the trabecular bone parameters or hematopoietic properties were not affected in mutant mice of either sex. The study uncovers a local source of IGF-1 in the bone marrow microenvironment that contributes to bone anabolic regulation in a site-specific manner.

**Lay Summary:** The insulin-like growth factor 1 (IGF-1) is known to play important roles in promoting bone growth and in regulating blood stem or progenitor cells. The factor is available both through the blood circulation and from the local production by cells in the bone tissue, but the functional contribution of each source is not well understood. Here we discover the non-blood cells in the bone marrow (bone marrow stromal cells) as a notable source of IGF-1 which specifically stimulates bone formation in the bone shaft of the long bone.

## Introduction

Insulin-like growth factor 1 (IGF-1), is among the most well characterized growth-promoting signals for both embryonic and postnatal tissues. In mouse knockout studies, IGF-1 deletion caused severe intrauterine growth retardation and perinatal mortality [1, 2]. Postnatally, IGF-1 functions both by mediating the function of growth hormone (GH) and through a GH-independent mechanism [3]. IGF-1 is abundantly produced by the liver and secreted to the circulation, but also generated in local tissues like bone [4]. In the serum, the vast majority of IGF-1 is bound to members of the IGF binding protein family (IGFBP 1-6), and the acid labile subunit (ALS) [5]. The association with ALS is believed to both prolong the half-life of IGFs and restrict its passage from the circulation to the extravascular compartment, thereby modulating its biological activities [6]. Evidence indicates that both circulating and locally produced IGF-1 contribute to growth regulation. In support of the endocrine function of hepatic IGF-1, liver-specific IGF-1 ablation combined with deletion of ALS resulted in >85% reduction in the circulating IGF-1 level and significant growth retardation and bone loss in postnatal mice [7]. On the other hand, liver-specific deletion of IGF-1 alone did not impair linear growth and only modestly reduced cortical bone growth even though the serum IGF-1 level was reduced by 75% [8–10]. The studies highlight a low threshold level of circulating IGF-1 necessary for supporting most of its endocrine function.

Genetic studies have identified important paracrine or autocrine functions for locally produced IGF-1 in promoting bone growth. Deletion of IGF-1 with either Col1a2-Cre in mesenchymal cells including osteoblasts, or Col2a1-Cre that targeted both chondrocytes and osteoblasts, suppressed skeletal growth and bone accrual without affecting serum IGF-1 levels [11, 12]. Deletion of IGF-1 in late osteoblasts and osteocytes with DMP1-Cre impaired both the normal growth of cortical bone and the anabolic response to mechanical loading [13, 14]. Conversely, IGF-1 overexpression from the osteocalcin promoter increased bone mineral density and bone formation rate [15]. Consistent with the ligand studies, deletion of the IGF-1 receptor (IGFR1) in osteoblasts with OC-Cre appeared to reduce bone formation by impeding mineralization, whereas deletion in preosteoblasts with Osx-Cre suppressed bone formation via impaired osteoblast differentiation [16, 17]. More recently, IGF-1 deletion in a subset of bone marrow stromal cells with LepR-Cre reduced both trabecular and cortical bone mass due to impaired bone formation [18]. Collectively, the studies establish that locally produced IGF-1 serves as an important anabolic signal for bone growth. Furthermore, the various local sources from different cell types likely contribute differently to the overall IGF-1 bioactivity in bone.

Local IGF-1 levels has been implicated in the regulation of hematopoiesis that occurs in the bone marrow microenvironment. A decline in bone marrow IGF-1 levels in middle-aged mice (∼12 months) has been shown to initiate hematopoietic stem cell (HSC) aging and induce myeloid-biased hematopoiesis [19]. However, as previous scRNA-seq studies have uncovered a high degree of heterogeneity among bone marrow stromal cells, it is important to delineate the cellular sources of IGF-1 responsible for osteogenic or hematopoietic activities in normal or pathological conditions [20, 21].

Here we study the physiological relevance of IGF-1 derived from the Adipoq-lineage bone marrow stromal cells. By deleting IGF-1 with Adipoq-Cre, we report cortical-specific bone loss without overt effects on hematopoietic stem and progenitor cells.

## Materials and Methods

### Mice

All mouse work was approved by the Children’s Hospital of Philadelphia Animal Care and Use Committee (IACUC approval number IAC#24-001296). Mice were housed at 22°C with a 12-hour light cycle (6 a.m. to 6 p.m.) and free access to food and water. Both male and female mice were analyzed in the study. Adipoq-Cre (Strain #: 028020) and IGF-1^f/f^ (Strain #: 016831) mouse lines were obtained from The Jackson Laboratory [22, 23].

### scRNA-seq analyses of bone marrow stromal cells (BMSC)

The scRNA-seq data was generated previously with FACS-purified endosteal stromal cells from tibias and femurs of 8-week-old male C57BL/6J mice [24] (Data accession GSE232738). scRNA-seq was performed with 10x Genomics. Downstream analyses were performed with Seurat 4.1.1 according to standard procedures.

### RT-qPCR in BMSC

IGF1 mRNA levels were measured by RT-qPCR in BMSC directly following isolation without culture. To isolate BMSC from the mice, the epiphyses were removed with scissors from femurs or tibias. The marrow content was flushed out with MEM-α media containing 10% FBS from both ends of the bone until the diaphysis appeared white. The cells were centrifuged and resuspended in MEM-α media containing 10% FBS and washed with PBS. Following this, the cells were mixed with CD45 microbeads (Miltenyi Biotec, 130-052-301), washed, centrifuged and resuspended before being passed though LD columns (Miltenyi Biotec, 130-042-901) on a MidiMACS™ separator (Miltenyi Biotec, 130-042-302) attached to a multistand. The flow-through CD45-negative cells were subjected to RNA extraction according to the RNeasy® Micro Kit (Qiagen). cDNA was made using the High-Capacity RNA-to-cDNA kit (Applied Biosystems). qPCR was performed with SYBR Green on QuantStudio3 (Applied Biosystems) with gene-specific primers for IGF-1 or HPRT (Table 1). Relative expression levels of IGF-1 were normalized to those of HPRT and calculated with the 2^-ΔΔCT^ method.

**Table 1.**
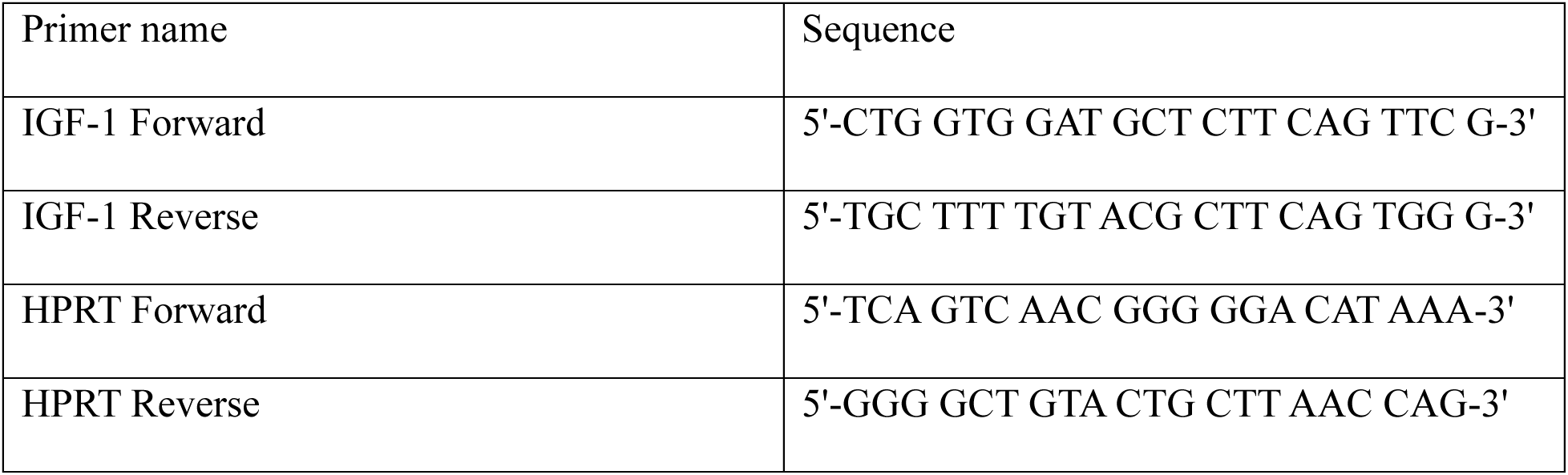
Sequence of qPCR Primers.

### Flow cytometry of BMSC

Targeting efficiency was assessed by tdTomato expression in central marrow versus endosteal BMSC from Adipoq-Cre; Ai9 mice at 2-3 months of age. After the epiphyses of tibias and femurs were excised with scissors and discarded, the central marrow fraction was collected by flushing with MEM-α media containing 10% FBS from both ends of the bone. To collect the endosteal fraction, the remaining bone was cut into small pieces and digested with 1 mg/ml dispase II (Roche, 4942078001) and 1 mg/ml STEMxyme1 (Worthington, LS004106) at 37 °C for 30 mins. The central marrow and endosteal fractions were cultured separately with MEM-α and 10% FBS for 7 days with media changed at day 4. The CD45^-^ BMSC were then purified from each culture with MACS as described above. Single cell suspensions were subjected to flow cytometry with CytoFLEX LX (Beckman Coulter) for tdTomato expression. Cells not expressing tdTomato were used as a gating control.

### µCT analyses

μCT analysis was performed with μCT45 (Scanco Medical AG, Switzerland) according to guidelines of the American Society of Bone and Mineral Research. The femurs were scanned with the X-ray source of 55 kVp, 145 µA, 8 W and 0.5 mm aluminum filter. The nominal voxel size was 4.5 μm and integration time was 400 ms. For quantifying trabecular bone parameters, two consecutive regions (ROI 1 and ROI 2) of 400 CT slices (1.8 mm) each were collected starting from 100 slices (0.45 mm) below the distal growth plate of the femur using a lower threshold of 350 and a Gaussian noise filter (sigma = 1.2, support = 2.0). For quantifying cortical bone parameters, 70 slices were analyzed using a lower threshold of 380 and a Gaussian noise filter (sigma = 1.2, support = 2.0). Males and females are quantified separately with genotypes blinded during analyses.

### Dynamic histomorphometry

For double labeling of bone forming surfaces, calcein (5 mg/mL, pH 7.2, 5μL/g body weight) (Sigma, C0875) was administered intraperitoneally seven days prior to harvest, followed by Alizarin Red S (15 mg/mL, pH 7.2, 5μL/g body weight) (Sigma, A5533) two days prior to harvest. Following harvest, femurs or tibias were fixed in 4% paraformaldehyde in PBS for 3 hrs at room temperature (RT) before being cryoprotected in 30% sucrose in PBS for 3 days at 4°C. The bones were then embedded in Tissue-Plus™ O.C.T. compound, cryosectioned at 10 μm using a cryostat (Leica CM1950) and adhered to cryofilm type II membrane (Section Lab, Co. Ltd.). Sections were washed in PBS and mounted with ProLong™ Gold Antifade Mountant (Thermo Fisher, P36930) before images were captured with ZEISS Axio Scan.Z1.

BIOQUANT® 2025 image analysis software was used for bone histomorphometry. Trabecular bone parameters were collected in ROI 1 starting at 450 µm below the growth plate with a fixed length of 1800 µm. The width of ROI 1 varied to encompass the entire trabecular bone width which differed across mice. The cortical region began adjacent to the end of ROI 1 at 2500 µm from the growth plate and extended for 3000 µm. Measurements included bone surface (mm), mineralizing surface (mm), bone formation rate (%), inter-label width (µm), mineral apposition rate (µm/day), and bone formation rate normalized by bone surface (µm/day). Genotypes of the mice were blinded during analyses.

### Serum PINP or CTX-1 ELISA

Whole blood was collected from mice via cardiac puncture using a 25 µm needle, and placed into an SST Microcontainer (BD) on ice. Tubes were centrifuged for 15 minutes at 4000g at 4°C to isolate serum. Serum samples were stored at -80 °C until use. PINP and CTX-1 were measured in diluted serum with Rat/Mouse PINP EIA and RatLaps® CTX-I EIA, respectively, by following the manufacturer’s instructions (Immuno Diagnostic Systems). The readings were acquired with a Cytation 5 image reader (Biotek).

### Immunofluorescence and quantification of bone marrow adipocytes

Femurs were fixed in 4% paraformaldehyde in PBS for 3 hrs at room temperature (RT) and then cryoprotected in 30% sucrose in PBS for 3 days at 4°C. They were embedded in Tissue-Plus™ O.C.T. compound, cryosectioned at 10 μm using a cryostat (Leica CM1950) and adhered to cryofilm type II membrane (Section Lab, Co. Ltd.).

For immunostaining, sections were washed in PBS for 15 mins, blocked with Antibody Diluent with BSA for 30 mins at RT, and incubated with rabbit anti-perilipin monoclonal antibody (1:200, Cell Signaling Technology, #9349) overnight at 4°C, followed by Alexa Fluor 647-conjugated F(ab’)2-goat anti-rabbit IgG (1:500, Thermo Fisher A21246) for 1 hr at RT. Sections were mounted with ProLong™ Gold Antifade Mountant (Thermo Fisher, P36930) and imaged with Leica TCS SP8 Confocal Microscope. Perilipin+ bone marrow adipocytes were quantified within an approximately 7 mm^2^ area encompassing the primary spongiosa and trabecular bone (3 mm in height) on three sections per femur.

### Analysis of bone marrow hematopoietic stem and progenitor cells

To isolate the bone marrow cells, both femurs were dissected from mice at P14 or 8 weeks of age. The femurs were cut at the metaphysis on both ends, and bone marrow was isolated by centrifugation using a quick pulse at 10,000 x g. Bone marrow was resuspended in PBS with 2% FBS. After cells were filtered through a 70μm filter, red cell lysis was performed to obtain bone marrow mononuclear cells. For flow cytometric analyses of HSPC cell surface markers, bone marrow mononuclear cells were incubated with a cocktail of anti-Ter119, B220, TCRβ, CD48, Gr-1, CD11b, c-kit, Sca-1, CD135, CD150, CD48 (BioLegend), and analyzed on a Cytek Aurora spectral flow cytometer with the FlowJo 10.10.0 software. HSPC populations were gated and quantified as previously described [25]. For colony forming assays, approximately 8000 bone marrow mononuclear cells were plated in fully supplemented methylcellulose (MethoCult GF M3434, Stemcell Technologies). Colonies were quantified after 7 days of culture based on morphology using phase contrast microscopy. All experiments included mice of both sexes.

## Results

### IGF-1 expression is enriched in CAR cells

To elucidate the molecular features of bone marrow stromal cells (BMSC), we have further analyzed the single-cell RNA-sequencing (scRNA-seq) dataset that we have previously generated [24]. Consistent with previous analyses, the predominant majority of BMSC (clusters 0, 1, 3-7) represents Cxcl12 abundant reticular (CAR) cells that are also enriched in Adipoq and LepR expression (Fig. 1A, B) [20, 21, 26]. Importantly, we discovered that IGF-1 was prominently expressed by most CAR cells (Fig. 1A, B). In particular, the CAR cells in clusters 0, 1, 3, 4 and 7 exhibited higher IGF-1 levels than osteoblasts (cluster 2) and the related cells (clusters 9, 11, 13) (Fig. 1A, B). As IGF-1 is known to function as an paracrine factor regulating bone formation and hematopoiesis, we decided to pursue the potential function of IGF-1 derived from the bone marrow CAR cells.

**Figure 1.**
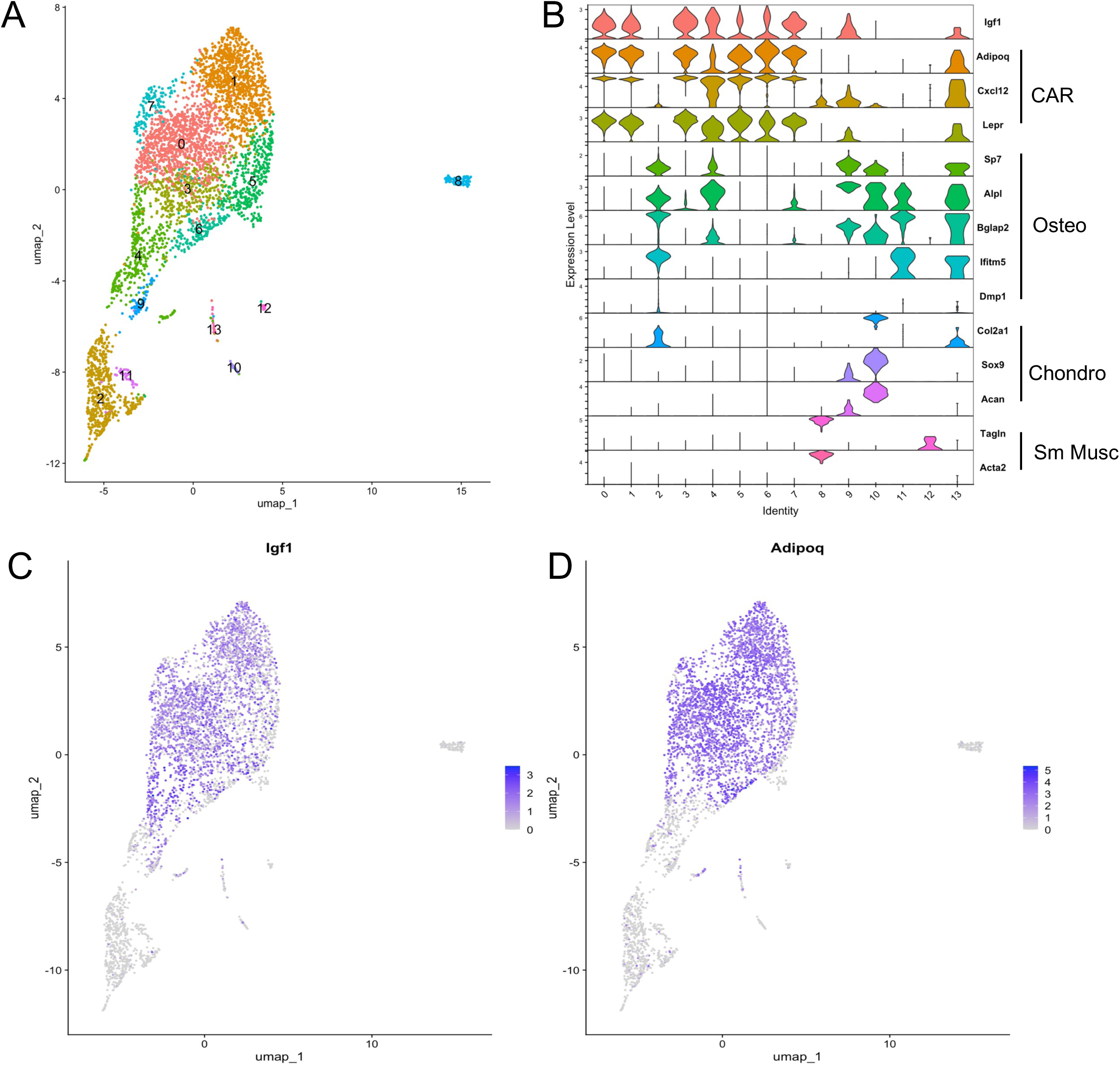
Enrichment of Igf1 mRNA among CAR cells in 8-week-old mice. (A) UMAP plot of bone marrow mesenchymal cell clusters by scRNA-seq. (B) Violin plots of select marker genes. CAR: Cxcl12-abundant reticular cells; Osteo: osteoblast; Chondro: chondrocyte; Sm Musc: Smooth Muscle. (C, D) Feature plots of Igf1 (C) and Adipoq (D) showing co-expression in CAR cells.

### Adipoq-Cre deletes IGF-1 in BMSC

The strong overlap between IGF-1 and Adipoq expression prompted us to employ Adipoq-Cre to delete IGF-1 in CAR cells. We first determined the efficacy of Adipoq-Cre in targeting BMSC in Adipoq-Cre;Ai9 mice. BMSC isolated from the central marrow versus endosteal niche were analyzed separately by flow cytometry. The results showed that ∼88% central marrow BMSC and ∼70% of endosteal BMSC expressed tdTomato, indicating adequate targeting of both populations by Adipoq-Cre (Fig. 2A-C). We next examined directly the deletion efficiency of IGF-1 in the BMSC of Adipoq-Cre;IGF-1^f/f^ mice (CKO). BMSC were enriched from flushed marrow through exclusion of the CD45^+^ cells with MACS (magnetic cell separation) beads, and then analyzed by RT-qPCR. The results showed notable reduction of IGF-1 mRNA in both male and female mice, indicating effective deletion of IGF-1 in BMSC by Adipoq-Cre (Fig. 2D, E). Thus, Adipoq-Cre provides a useful tool for restricting IGF-1 production by BMSC.

**Figure 2.**
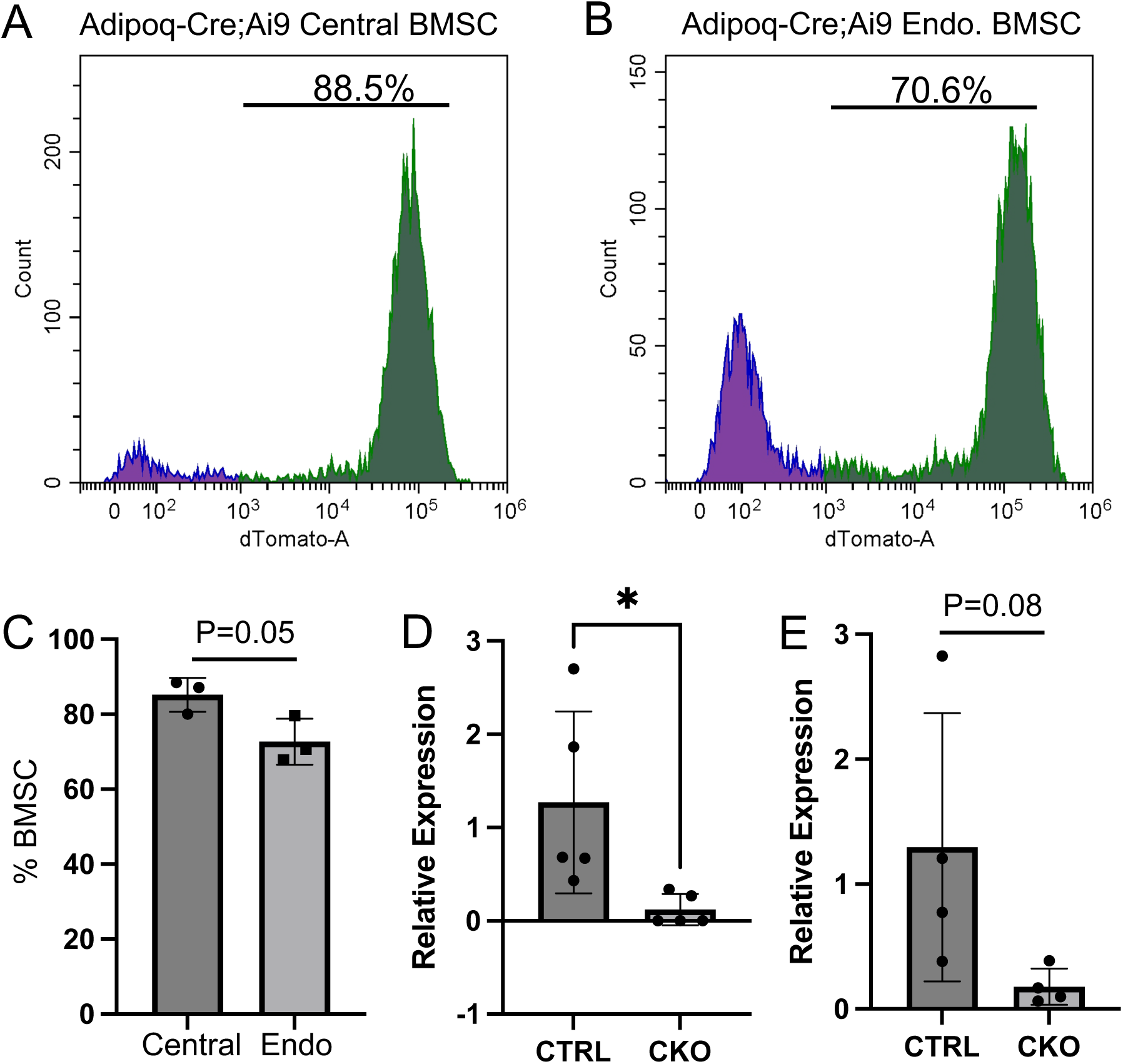
Targeting of BMSC by Adipoq-Cre. (A, B) Representative flow cytometry graphs showing tdTomato detection in central marrow (A) versus endosteal fraction (B) of BMSC from a 9-week-old Adipoq-Cre;Ai9 female mouse. (C) Quantification of targeting efficiency in central marrow versus endosteal BMSC in Adipoq-Cre;Ai9 female mice at 9-12 weeks of age. (D, E) Relative levels of IGF-1 mRNA in BMSC from female (D) or male (E) mice at 8 weeks of age. CTRL: Igf1^f/f^; CKO: Adipoq-Cre;Igf1^f/f^. Statistics: unpaired t test, *p <0.05, each dot representing a single mouse.

### IGF-1 deletion causes age-dependent defect in endocortical bone formation in female mice

We next examined the potential effects of IGF-1 deletion on bone parameters by µCT. We evaluated the cortical bone at the mid-diaphysis and the trabecular bone in upper (ROI 1) versus lower (ROI 2) regions. To our surprise, at 8 weeks of age, no abnormalities in any of the cortical or trabecular bone parameters were observed in Adipoq-Cre;IGF-1^f/f^ (CKO) mice of either sex (Supplemental Fig. S1, S2). As a previous study reported that deletion of IGF-1 by LepR-Cre reduced bone mass in 12-week-old mice, we suspected that BMSC-derived IGF-1 might influence bone mass in an age-specific manner [18]. To test this notion, we analyzed the bones of female CKO versus control (CTRL) mice at 12 weeks of age. Here, like in the younger mice, all trabecular bone parameters were normal in the CKO mice (Fig. 3A, B). However, notable defects were detected in the cortical bone, as indicated by significant reductions in bone area (BA), bone area fraction (BA/TA) and cortical thickness (C. Th), without changes in the total cross-sectional area (TA) (Fig. 3C, D). Thus, whereas much of the BMSC-derived IGF-1 appears to be dispensable for trabecular bone mass or organization, it is necessary to support a normal cortical bone thickness during the third month of postnatal life when bone continues to grow in mice.

**Figure 3.**
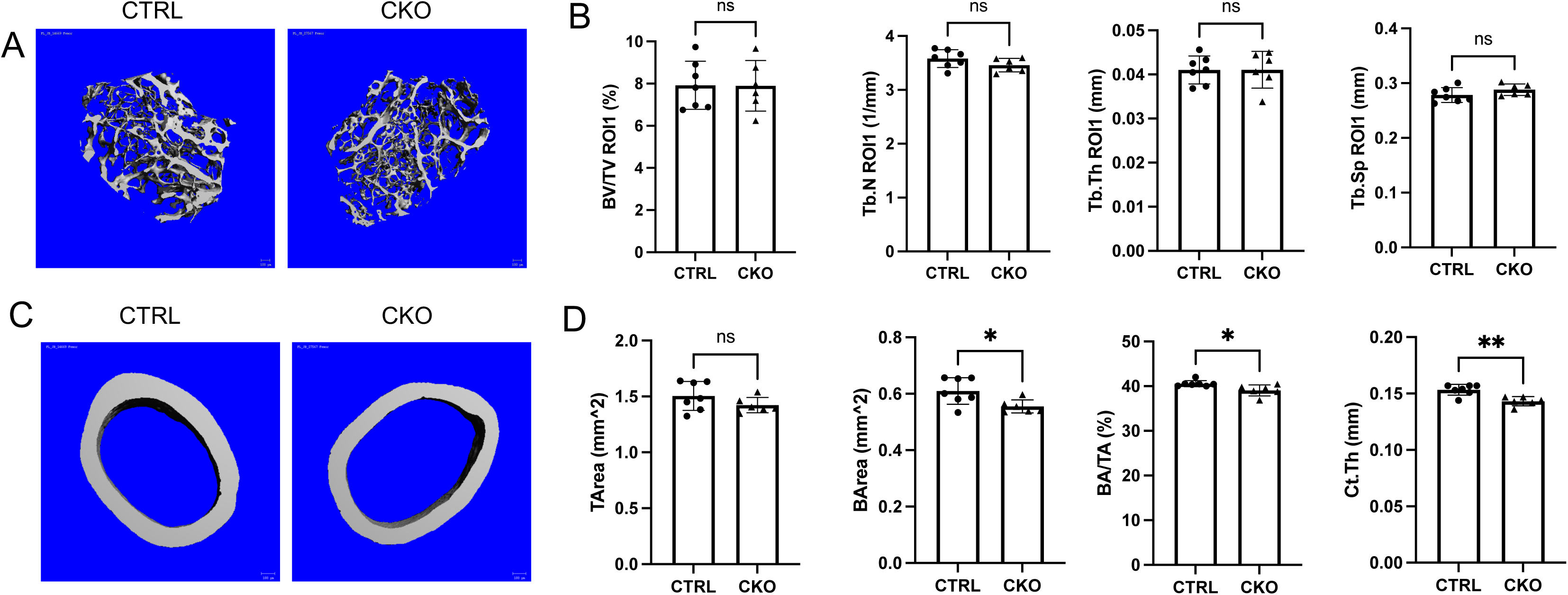
Effects of BMSC-derived IGF-1 on bone mass in 12-week-old female mice. (A, C) Representative µCT 3-D reconstruction images of trabecular (A) or cortical (C) bone. (B, D) Quantification of trabecular (B) versus cortical (D) bone parameters by µCT analyses. CTRL: Igf1^f/f^; CKO: Adipoq-Cre;Igf1^f/f^. Statistics: unpaired t test, *p<0.05, **p<0.01, ns: non-significant, p>0.05, each dot representing a single mouse.

We next assessed the cellular basis for the reduced cortical bone thickness in the 12-week-old female CKO mice. Dynamic histomorphometry showed that bone formation rate (BFR/BS) at the endosteal bone surface was significantly reduced owing to suppressed mineral apposition rate (MAR) even though mineralizing surface areas (MS/BS) were not altered (Fig. 4A). In contrast, the trabecular bone formation parameters were normal in the CKO mice (Fig. 4B). Finally, serum biochemistry did not reveal any significant changes in the overall bone formation (P1NP) or bone resorption (CTX-I) activity in the CKO versus CTRL mice (Fig. 5A, B). Thus, site-specific impairment in bone formation at the endosteal bone surface contributes to the cortical bone defect upon IGF-1 deletion in BMSC.

**Figure 4.**
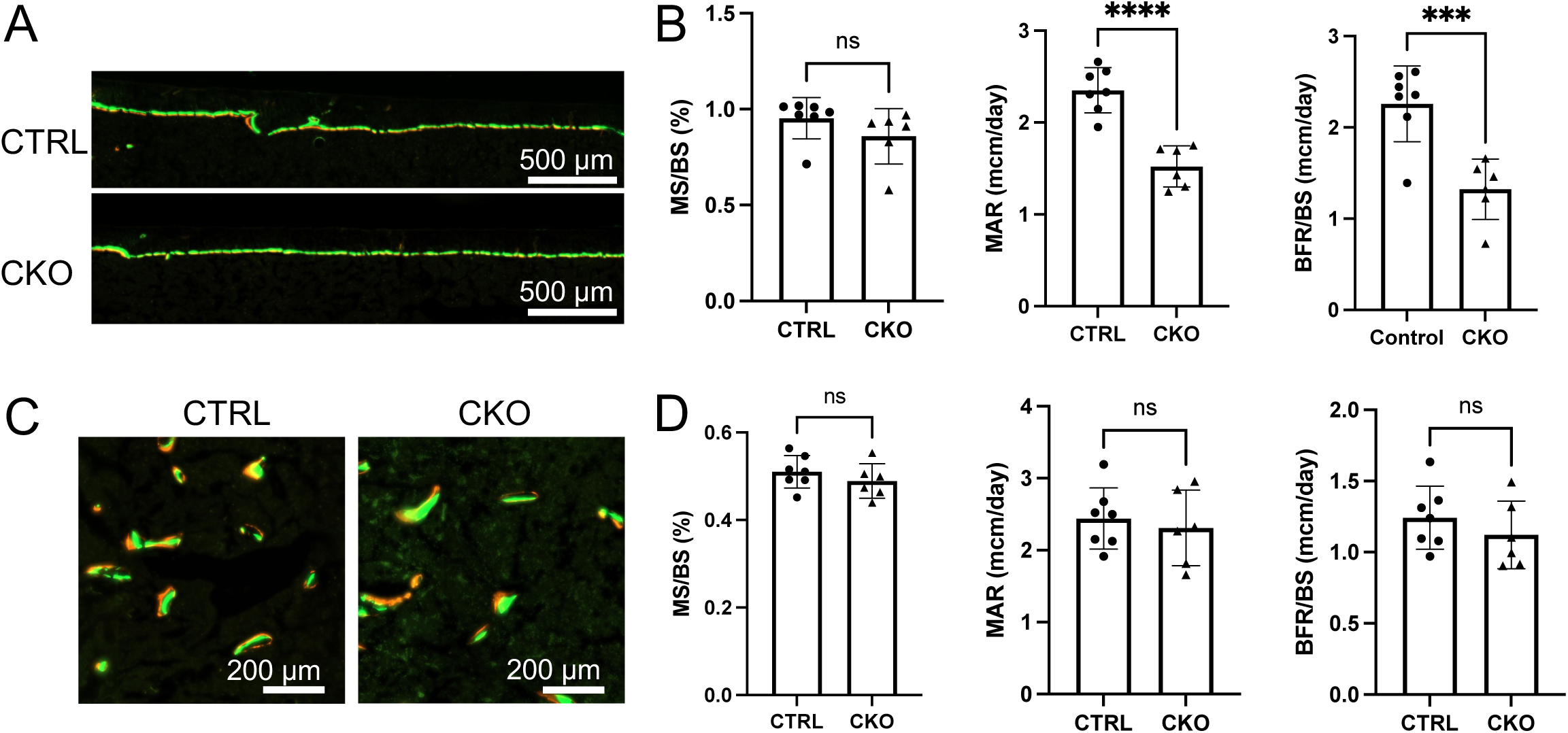
Dynamic histomorphometry for bone formation in 12-week-old female mice. (A) Representative images of endosteal bone labeling. (B) Parameters for endosteal bone surface. (C) Representative images of trabecular bone labeling. (D) Parameters for trabecular bone surface. MS: mineralizing surface; BS: bone surface; MAR: mineral apposition rate; BFR: bone formation rate. CTRL: Igf1^f/f^; CKO: Adipoq-Cre;Igf1^f/f^. Statistics: unpaired t test, ***p<0.001, ****p<0.0001, ns: non-significant, p>0.05, each dot representing a single mouse.

**Figure 5.**
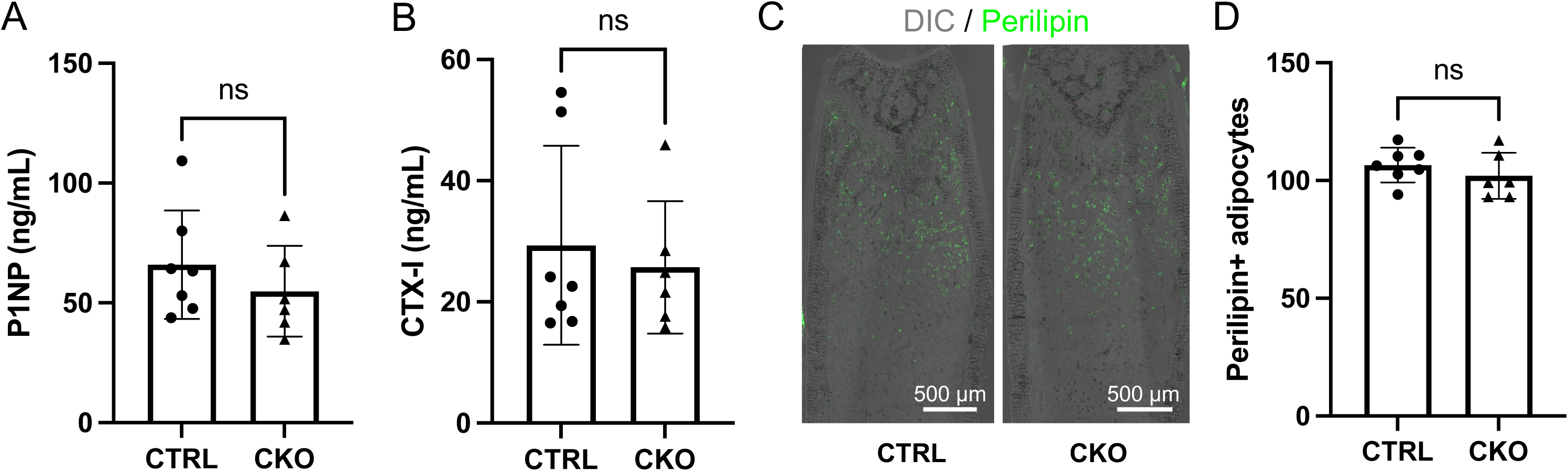
Quantification of serum bone markers and marrow adiposity. (A, B) Serum levels of bone formation (A) and resorption (B) markers. (C) Representative images for immunofluorescence staining of perilipin 1 (green) on sections of distal femur. Tissue morphology shown by DIC (differential interference contrast) imaging. (D) Quantification of adipocyte numbers in distal femur. CTRL: Igf1^f/f^; CKO: Adipoq-Cre;Igf1^f/f^. Statistics: unpaired t test, ns: non-significant, p>0.05, each dot representing a single mouse.

IGF-1 is well established to promote adipocyte differentiation from preadipocytes [27]. We therefore examined the potential effect of IGF-1 deletion on bone marrow adiposity in the CKO mice. Immunofluorescence staining with a perilipin 1 antibody revealed normal adipocyte numbers in the long bones of the mutant mice (Fig. 5C, D). Thus, IGF-1 originated from the Adipoq-lineage BMSC is likely dispensable for bone marrow adipogenesis under normal conditions.

### IGF-1 from Adipoq-lineage BMSC is dispensable for hematopoiesis

We next determined whether loss of IGF-1 in the bone marrow microenvironment impaired hematopoietic stem and progenitor cells (HSPC). For this, we isolated bone marrow cells from the femurs at either 2 or 8 weeks of age, and performed flow cytometry and methylcellulose colony forming assays with the bone marrow mononuclear cells. Flow cytometry indicated that the fractions of total Lin^-^Sca1^+^c-Kit^+^ (LSK) cells, long-term or short-term HSCs, and various multipotent progenitors (MPP2-4) were all normal among the bone marrow mononuclear cells in the CKO mice of either age (Fig. 6A, B). MPP designations were based on a previous publication [25]. Moreover, the colony forming assays revealed no effect on the multipotential myeloid progenitors (CFU-GEMM), erythroid burst-forming units (BFU-E) or granulocyte-macrophage progenitors (CFU-G/M/GM) by IGF-1 deletion (Fig. 6C). Thus, IGF-1 production by Adipoq-lineage BMSC appears to have no discernible effect on HSPC population abundance or colony-forming potential in young postnatal mice.

**Fig. 6.**
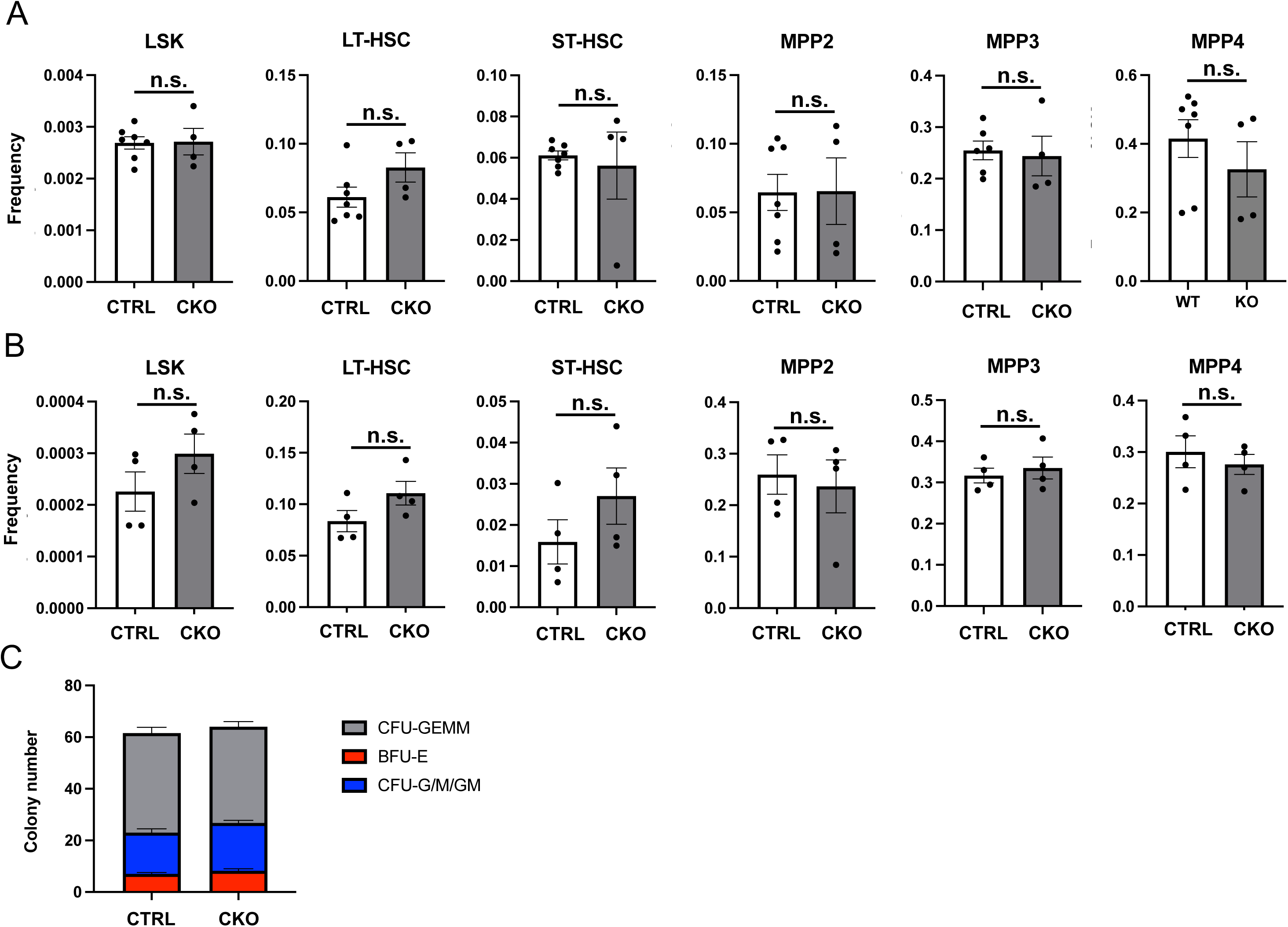
Quantification of bone marrow HSPC populations and colony formation potential. (A-B) Analysis of HSPC populations by flow cytometry in mice at 2 (A) or 8 (B) weeks of age. HSPC populations expressed as absolute frequencies among bone marrow mononuclear cells. (C) Methylcellulose colony forming assays of bone marrow mononuclear cells derived from 2-wk-old mice. Colonies identified by morphology and counted at day 7 (n = 4 per genotype). CTRL: *Igf1^f/f^*; CKO: *AdipoqCre^+^ Igf1^f/f^*. Acronyms: LSK, Lineage^-^Sca1^+^c-Kit^+^ hematopoietic stem and progenitor cells; LT-HSC, long-term hematopoietic stem cell; ST-HSC, short-term hematopoietic stem cell; MPP2, multipotent progenitor population 2 (megakaryocyte and erythroid bias); MMP3, multipotent progenitor population 3 (granulocyte and macrophage bias); MPP4, multipotent progenitor population 4 (lymphoid bias); CFU-GEMM, colony forming unit – Granulocyte, Erythrocyte, Megakaryocyte, Monocyte; BFU-E, burst forming unit – Erythrocyte; CFU-G/M/GM, colony forming unit – Granulocyte, Macrophage. Data represented as mean ± SEM. Statistics: unpaired t-tests; n.s. (non-significant): p>0.05, each dot representing a single mouse in A and B.

## Discussion

We have investigated the potential paracrine function of IGF-1 derived from BMSC in the regulation of bone growth and hematopoiesis. By deleting IGF-1 with Adipoq-Cre which targets most BMSC, the study uncovered a site-specific contribution of the local IGF-1 to bone formation at the endosteum of long bones, resulting in thinner cortices by 12 weeks of age. On the other hand, the trabecular bone parameters or the HSPC frequencies were not affected by the deletion. Together with previous studies, the current findings support the view that multiple sources of IGF-1 in the bone marrow environment likely exert distinct niche functions towards osteogenic or hematopoietic cells.

As Adipoq-Cre also targets mature adipocytes, it is important to note that a previous study reported no changes to adipose depots, whole-body metabolism or circulating IGF-1 levels when IGF-1 was deleted with Adipoq-Cre in mice under normal feeding conditions [28]. Thus, the endocortical bone phenotype observed in the mutant mice here is most consistent with the paracrine function of IGF-1 locally produced by BMSC.

We observed no HSPC phenotype in the CKO mice. This is consistent with a previous study where deletion of IGF-1 in BMSC with LepR-Cre did not impair hematopoiesis [18]. In contrast, IGF-1 deletion with Nestin-Cre^ER^ was reported to cause HSC aging and myeloid-biased hematopoiesis [29], or impaired bone formation [30]. However, a similar Nestin-Cre^ERT2^ line was found to target mostly endothelial cells (a known source of IGF-1) instead of stromal cells in the bone marrow of postnatal developing mice [31]. Thus, it remains to be further determined whether the phenotypes reported earlier were strictly dependent on IGF-1 produced by BMSC.

The bone phenotype here seems to be at odds with a previous report that deletion of IGF-1 with LepR-Cre significantly reduced both trabecular and cortical bone mass due to impaired bone formation [18]. The LepR-Cre-mediated deletion also increased bone marrow fat which was not observed in the current study. Although LepR-Cre differs from Adipoq-Cre with additional activity in the periosteum, they both target BMSC with high efficiency. It is therefore surprising that deletion of IGF-1 by each Cre resulted in seemingly different bone phenotypes. However, some technical differences should be noted. Whereas the previous study focused on male mice at a single time point (12 weeks of age), we analyzed both sexes at 8 weeks and only the females at 12 weeks of age. Whether the age and/or sex differences could explain some of the phenotypic discrepancy is not clear at present. A more interesting alternative is that the two Cre drivers may target subsets of BMSC with different efficiencies. If so, the different phenotypes would highlight distinct contributions of IGF-1 from the various BMSC sources to trabecular versus endocortical bone formation. Future studies are necessary to examine potential functional diversity across subpopulations of BMSC.

## Acknowledgements

The work is partially supported by NIH grants R01 AG077911 (FL), R01 DK125498 (FL) and NHLBI K99 HL156052 (CST).

## Supplemental Figure Legends

**Figure S1.**
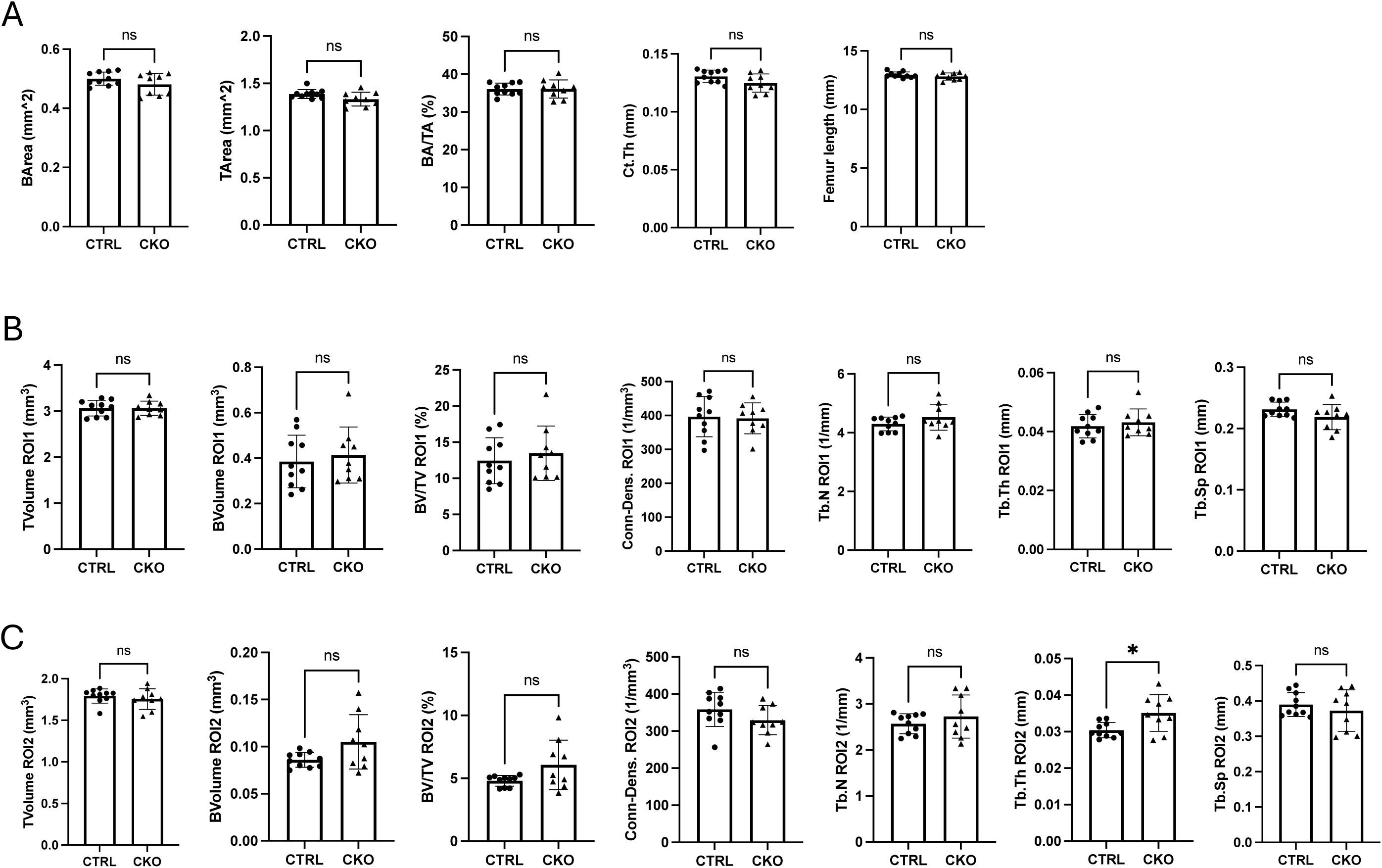
µCT quantification of bone parameters in the femur of 8-week-old female mice. (A) Cortical bone parameters. BA: bone area; TA: tissue area; Ct. Th: cortical thickness (B, C) Trabecular bone parameters in region of interest 1 (ROI 1) (B) and ROI 2 (C). BV: bone volume; TV: total volume; Conn. Dens.: connectivity density; Tb. N. trabeculae number; Tb. Th: trabecular thickness; Tb. Sp.: trabecular spacing. *: unpaired t test, p<0.05, error bar: SD, each dot represents an individual mouse. CTRL: Igf1^f/f^; CKO: Adipoq-Cre;Igf1^f/f^.

**Figure S2.**
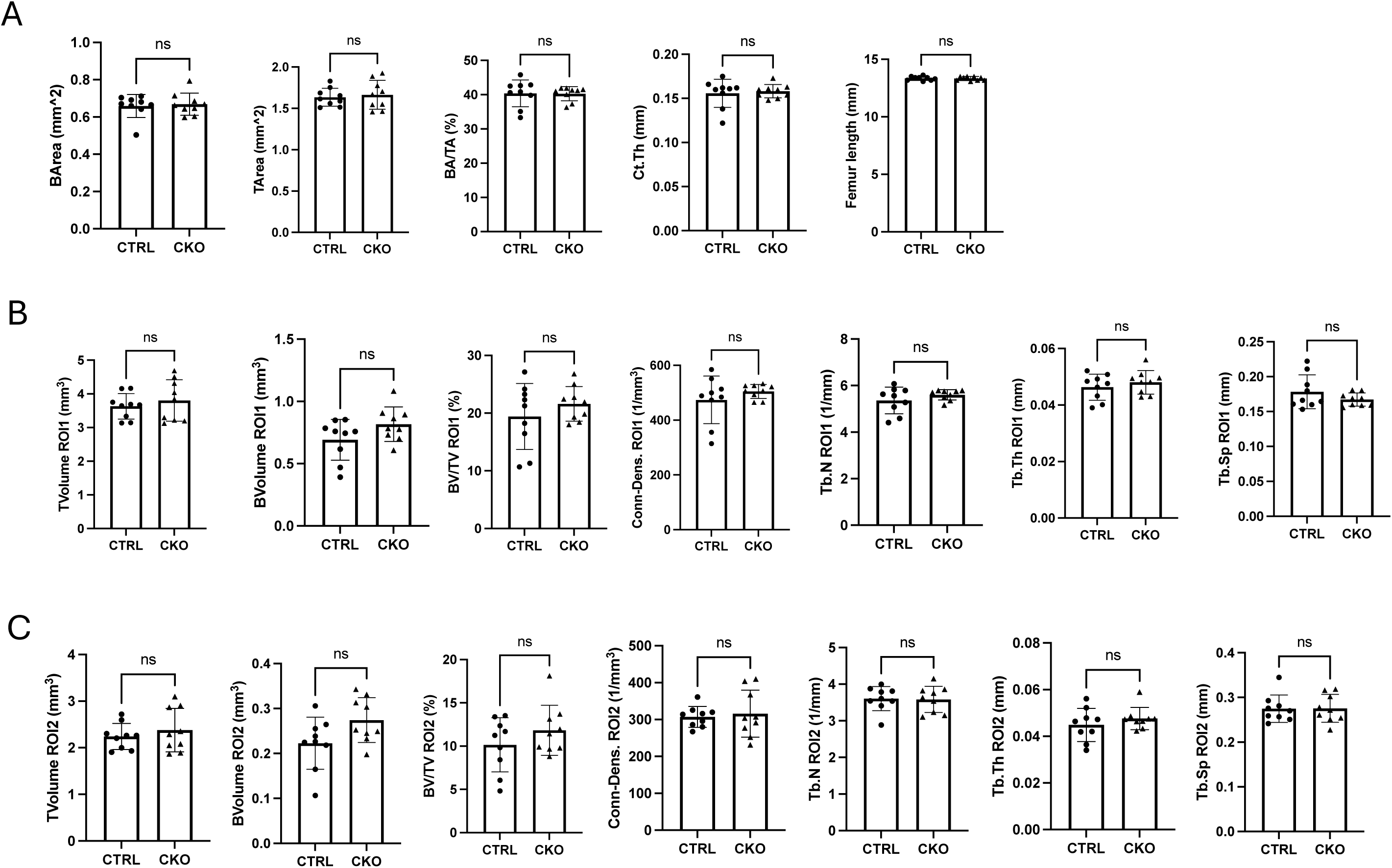
µCT quantification of bone parameters in the femur of 8-week-old male mice. (A) Cortical bone parameters. BA: bone area; TA: tissue area; Ct. Th: cortical thickness (B, C) Trabecular bone parameters in region of interest 1 (ROI 1) (B) and ROI 2 (C). BV: bone volume; TV: total volume; Conn. Dens.: connectivity density; Tb. N. trabeculae number; Tb. Th: trabecular thickness; Tb. Sp.: trabecular spacing. Statistics: unpaired t test, * p<0.05, ns (non-significant) p>0.05, error bar: SD, each dot represents an individual mouse. CTRL: Igf1^f/f^; CKO: Adipoq-Cre;Igf1^f/f^.

**Figure S3.**
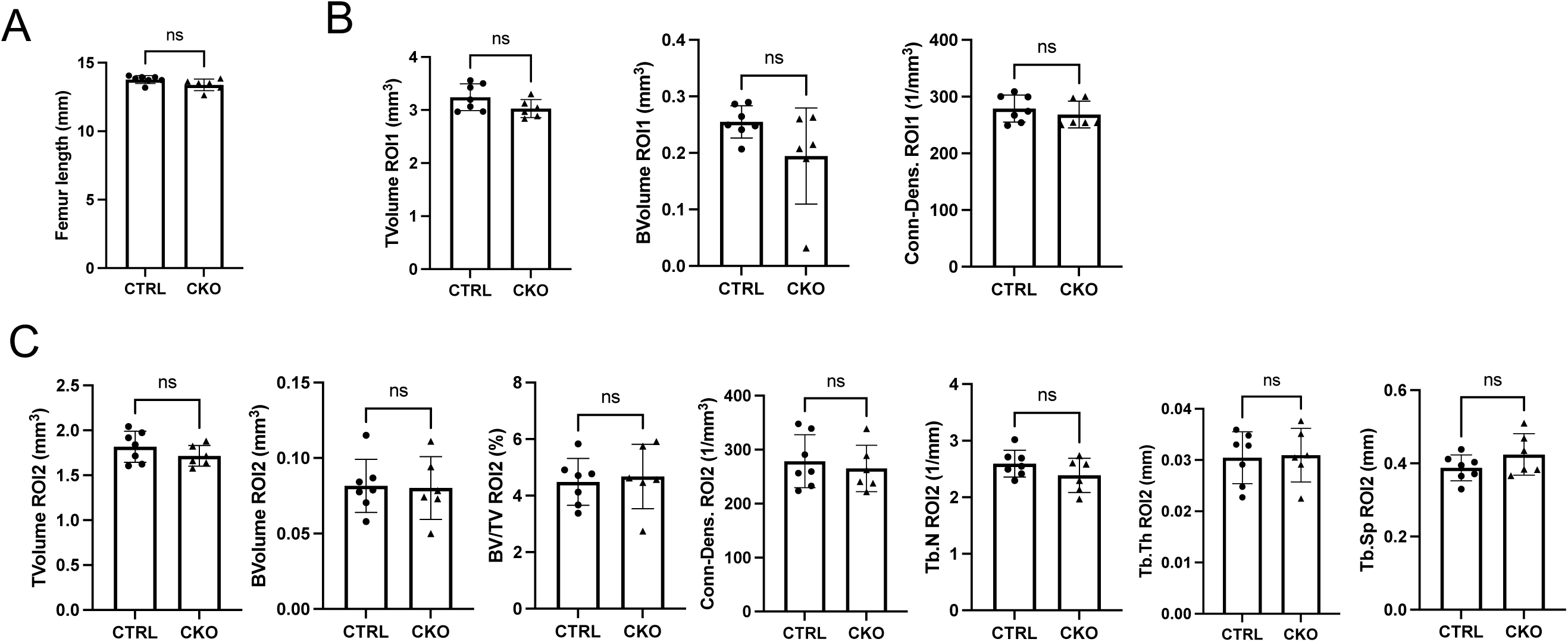
µCT quantification of bone parameters in the femur of 12-week-old female mice. (A) Femur lengths. (B, C) Trabecular bone parameters in region of interest 1 (ROI 1) (B) and ROI 2 (C). BV: bone volume; TV: total volume; Conn. Dens.: connectivity density; Tb. N. trabeculae number; Tb. Th: trabecular thickness; Tb. Sp.: trabecular spacing. Statistics: unpaired t test, ns: p>0.05, error bar: SD, each dot represents an individual mouse. CTRL: Igf1^f/f^; CKO: Adipoq-Cre;Igf1^f/f^.

## References

1. Liu, J.P., et al., Mice carrying null mutations of the genes encoding insulin-like growth factor I (Igf-1) and type 1 IGF receptor (Igf1r). Cell, 1993. 75(1): p. 59–72.

2. Baker, J., et al., Role of insulin-like growth factors in embryonic and postnatal growth. Cell, 1993. 75(1): p. 73–82.

3. Lupu, F., et al., Roles of growth hormone and insulin-like growth factor 1 in mouse postnatal growth. Dev Biol, 2001. 229(1): p. 141–62.

4. Kawai, M. and C.J. Rosen, The insulin-like growth factor system in bone: basic and clinical implications. Endocrinol Metab Clin North Am, 2012. 41(2): p. 323–33, vi.

5. Rechler, M.M. and D.R. Clemmons, Regulatory Actions of Insulin-like Growth Factor-binding Proteins. Trends in endocrinology and metabolism: TEM, 1998. 9(5): p. 176–83.

6. Boisclair, Y.R., et al., The acid-labile subunit (ALS) of the 150 kDa IGF-binding protein complex: an important but forgotten component of the circulating IGF system. J Endocrinol, 2001. 170(1): p. 63–70.

7. Yakar, S., et al., Circulating levels of IGF-1 directly regulate bone growth and density. J Clin Invest, 2002. 110(6): p. 771–81.

8. Sjogren, K., et al., Liver-derived insulin-like growth factor I (IGF-I) is the principal source of IGF-I in blood but is not required for postnatal body growth in mice. Proc Natl Acad Sci U S A, 1999. 96(12): p. 7088–92.

9. Yakar, S., et al., Normal growth and development in the absence of hepatic insulin-like growth factor I. Proc Natl Acad Sci U S A, 1999. 96(13): p. 7324–9.

10. Yakar, S., et al., Serum IGF-1 determines skeletal strength by regulating subperiosteal expansion and trait interactions. J Bone Miner Res, 2009. 24(8): p. 1481–92.

11. Govoni, K.E., et al., Conditional deletion of insulin-like growth factor-I in collagen type 1alpha2-expressing cells results in postnatal lethality and a dramatic reduction in bone accretion. Endocrinology, 2007. 148(12): p. 5706–15.

12. Govoni, K.E., et al., Disruption of insulin-like growth factor-I expression in type IIalphaI collagen-expressing cells reduces bone length and width in mice. Physiol Genomics, 2007. 30(3): p. 354–62.

13. Sheng, M.H., et al., Disruption of the insulin-like growth factor-1 gene in osteocytes impairs developmental bone growth in mice. Bone, 2013. 52(1): p. 133–44.

14. Lau, K.H., et al., Osteocyte-derived insulin-like growth factor I is essential for determining bone mechanosensitivity. Am J Physiol Endocrinol Metab, 2013. 305(2): p. E271–81.

15. Zhao, G., et al., Targeted Overexpression of Insulin-Like Growth Factor I to Osteoblasts of Transgenic Mice: Increased Trabecular Bone Volume without Increased Osteoblast Proliferation*. Endocrinology, 2000. 141(7): p. 2674–2682.

16. Zhang, M., et al., Osteoblast-specific knockout of the insulin-like growth factor (IGF) receptor gene reveals an essential role of IGF signaling in bone matrix mineralization. J Biol Chem, 2002. 277(46): p. 44005–12.

17. Xian, L., et al., Matrix IGF-1 maintains bone mass by activation of mTOR in mesenchymal stem cells. Nature medicine, 2012. 18(7): p. 1095–101.

18. Wang, J., et al., Bone marrow-derived IGF-1 orchestrates maintenance and regeneration of the adult skeleton. Proc Natl Acad Sci U S A, 2023. 120(1): p. e2203779120.

19. Young, K., et al., Decline in IGF1 in the bone marrow microenvironment initiates hematopoietic stem cell aging. Cell Stem Cell, 2021. 28(8): p. 1473–1482.e7.

20. Baccin, C., et al., Combined single-cell and spatial transcriptomics reveal the molecular, cellular and spatial bone marrow niche organization. Nat Cell Biol, 2020. 22(1): p. 38–48.

21. Zhong, L., et al., Single cell transcriptomics identifies a unique adipose lineage cell population that regulates bone marrow environment. Elife, 2020. 9.

22. Eguchi, J., et al., Transcriptional control of adipose lipid handling by IRF4. Cell Metab, 2011. 13(3): p. 249–59.

23. Liu, J.L., et al., Insulin-like growth factor-I aXects perinatal lethality and postnatal development in a gene dosage-dependent manner: manipulation using the Cre/loxP system in transgenic mice. Mol Endocrinol, 1998. 12(9): p. 1452–62.

24. Ji, X., et al., Genetic activation of glycolysis in osteoblasts preserves bone mass in type I diabetes. Cell Chem Biol, 2023. 30(9): p. 1053–1063 e5.

25. Hall, T.D., et al., Murine fetal bone marrow does not support functional hematopoietic stem and progenitor cells until birth. Nat Commun, 2022. 13(1): p. 5403.

26. Sugiyama, T., et al., Maintenance of the hematopoietic stem cell pool by CXCL12-CXCR4 chemokine signaling in bone marrow stromal cell niches. Immunity, 2006. 25(6): p. 977–88.

27. Tang, Q.Q. and M.D. Lane, Adipogenesis: from stem cell to adipocyte. Annu Rev Biochem, 2012. 81: p. 715–36.

28. Chang, H.R., et al., Macrophage and adipocyte IGF1 maintain adipose tissue homeostasis during metabolic stresses. Obesity (Silver Spring), 2016. 24(1): p. 172–83.

29. Young, K., et al., Decline in IGF1 in the bone marrow microenvironment initiates hematopoietic stem cell aging. Cell Stem Cell, 2021. 28(8): p. 1473–1482 e7.

30. Crane, J.L., et al., IGF-1 Signaling is Essential for DiXerentiation of Mesenchymal Stem Cells for Peak Bone Mass. Bone Res, 2013. 1(2): p. 186–94.

31. Ono, N., et al., Vasculature-associated cells expressing nestin in developing bones encompass early cells in the osteoblast and endothelial lineage. Dev Cell, 2014. 29(3): p. 330–9.

